# Entorhinal grid coding as a functional link between tau accumulation and episodic memory in human aging

**DOI:** 10.64898/2026.08.26.747192

**Authors:** Vladislava Segen, Tuğçe Belge Bernard, Gerard Callau Navarro, Philip Bahrd, Niklas Behrenbruch, Beate Schumann-Werner, Svenja Schwarck, Berta Garcia-Garcia, Henryk Barthel, Osama Sabri, Michael C. Kreissl, Emrah Düzel, Anne Maass, Thomas Wolbers

## Abstract

Episodic memory decline is a common feature of cognitively normal aging, but its extent varies markedly across individuals. Although entorhinal tau pathology is thought to be a key contributor to episodic memory impairment, the neural mechanisms linking early tau accumulation to memory differences remain unclear. Grid-cell computations in the entorhinal cortex, which provide scaffolds for organizing experiences into episodic memories, offer one candidate mechanism. Here, we combined virtual-reality functional MRI, multivariate analysis, tau PET, and delayed word-list recall in cognitively normal older adults to test whether tau-related alterations in entorhinal coding are associated with worse episodic memory. Weaker left entorhinal grid-cell-like signal was associated with poorer memory performance, and individuals with higher left entorhinal tau burden showed weaker grid-cell-like signal. This association was specific to the canonical six-fold signal and was not explained by entorhinal volume, mean diffusivity, or intracortical myelination. A cross-sectional Bayesian mediation analysis further demonstrated that bilateral medial temporal tau burden is related to memory indirectly through left entorhinal grid-cell-like signal. Together, these findings provide evidence that entorhinal grid codes may constitute a functional pathway linking tau accumulation to memory variability in normal aging.

## Introduction

Cognitive aging is associated with a decline in episodic memory^1^, with consequences that can affect everyday functioning, independence, and quality of life. Episodic memory relies on entorhinal-hippocampal codes that bind event content with spatial and temporal context into unique memory representations^2–5^. Age-related changes in entorhinal–hippocampal coding may therefore contribute to individual differences in memory performance among older adults^6–8^. For example, cognitively normal older adults show reduced anterolateral entorhinal engagement together with increased dentate gyrus/CA3 activity during memory discrimination tasks, and this imbalance has been linked to poorer memory performance^9^. Within this circuit, the entorhinal cortex plays a central role in interfacing hippocampal memory representations with cortical information about objects, space, and context^10,11^.

One candidate computation that may contribute to age-related memory differences is provided by entorhinal grid cells, which are thought to supply a stable spatial metric for anchoring experiences to where they occurred^2^, thereby contributing to the spatial scaffold of episodic memory. Beyond navigation, entorhinal grid-cell-like activity has been observed during conceptual knowledge representation^12–14^, including representations in which it reflects the relational structure of newly learned word meaning^13^. Computational work has further proposed that grid-cell firing may arise during memory encoding and retrieval of spatial and non-spatial attributes^2,15^. Together, this theoretical, empirical, and computational work suggests that grid-cell-like representations provide a core entorhinal scaffold for organizing experiences into episodic memories.

If grid-cell-like codes provide a scaffold for memory representations, then age-related changes in these codes may contribute to differences in memory performance among older adults. Indeed, rodent work indicates that aging alters grid-cell coding properties: aged animals show reduced spatial firing stability despite largely preserved grid-cell density^16^. Similar findings have been observed in the hippocampus, where aged rats show reduced temporal stability of place-cell firing without an apparent reduction in the absolute number of place cells^17^. Human fMRI evidence points in a similar direction. Older adults show reduced entorhinal grid-cell-like representations, while preserved grid-cell-like coding in older adults is associated with better spatial navigation performance^18^. Together, this work indicates that entorhinal grid-cell-like codes are altered in aging; however, it remains unknown whether such alterations also contribute to age-related variability in episodic memory.

A second knowledge gap pertains to the neurophysiological mechanisms that might underlie this age-related disruption in grid-cell-like codes. One candidate is the accumulation of hyperphosphorylated tau, an abnormally modified form of the microtubule-associated protein tau that is best known for its role in Alzheimer’s disease pathology. Importantly, tau can also accumulate during cognitively normal aging^19,20^, and it has been linked to episodic memory decline^20–23^. Human PET and histology studies show that tau pathology emerges earliest in the entorhinal cortex and then spreads through the medial temporal lobe^20,24,25^, positioning the circuitry that supports grid-cell computations at the frontline of age-related tau accumulation. Consistent with this vulnerability, rodent work has demonstrated that early entorhinal tau pathology is sufficient to compromise grid-cell coding, alongside impairments in spatial memory^26–28^. Together, these findings suggest that tau pathology may progressively degrade the entorhinal computations required for grid-cell coding.

Our lack of understanding of how early molecular pathology may translate into computational dysfunction and, ultimately, episodic memory decline in older adults is a key obstacle for the development of mechanistically grounded interventions^29^. To address these knowledge gaps, we combined second generation[¹⁸F]PI-2620 tau PET imaging with functional MRI during spatial navigation and delayed word-list recall to test whether (1) entorhinal grid-cell-like representations relate to episodic memory performance; (2) early tau pathology in normal aging is associated with altered entorhinal grid-cell-like representations; and (3) tau-related disruption of this entorhinal computation may provide a mechanistic pathway linking tau pathology to episodic memory decline. To preview, our results provide converging evidence that early tau pathology in normal aging is linked to disrupted grid-cell-like representations, and that these representations mediate the relationship between age-related tau accumulation and episodic memory performance. By identifying altered entorhinal grid-cell-like codes as a potential mechanistic link between tau pathology and memory decline, this work identifies a potential targetable functional mechanism for future diagnostic tools and interventions aimed at preserving cognitive health in aging.

## Results

### Weaker left entorhinal grid-cell-like signal is associated with lower episodic memory performance in healthy older adults

We first asked whether entorhinal grid-cell-like coding was related to episodic memory performance in 93 cognitively normal older adults (see Methods). To do this, participants completed the German version of the Rey Auditory Verbal Learning Test^30^ (RAVLT), a standard word-list learning task with a delayed recall used to assess verbal episodic memory (Fig 1a). Participants also completed a virtual-reality spatial navigation task during fMRI in which they passively navigated through a square environment containing five unique objects and were instructed to remember each object’s location (Fig. 1b). These behavioural and fMRI data were subsequently combined with tau PET and MRI-derived measures of entorhinal structure and tissue integrity, providing the multimodal entorhinal dataset analysed throughout the study (Fig. 1c). The task was performed across two runs, with a different set of objects in each run. Participants performed above chance on the three-alternative forced-choice object-location test following navigation, indicating that they attended to the virtual environment and encoded object locations (mean accuracy across runs = 0.55, SD = 0.22; chance level = 0.33; one-sample test against chance: p <0.001). To quantify grid-cell-like representations, we used a multivariate representational similarity analysis (see Methods and Fig. 2a). This allowed us to test whether individual differences in entorhinal grid-cell-like signal were associated with episodic memory performance.

**Fig. 1.**
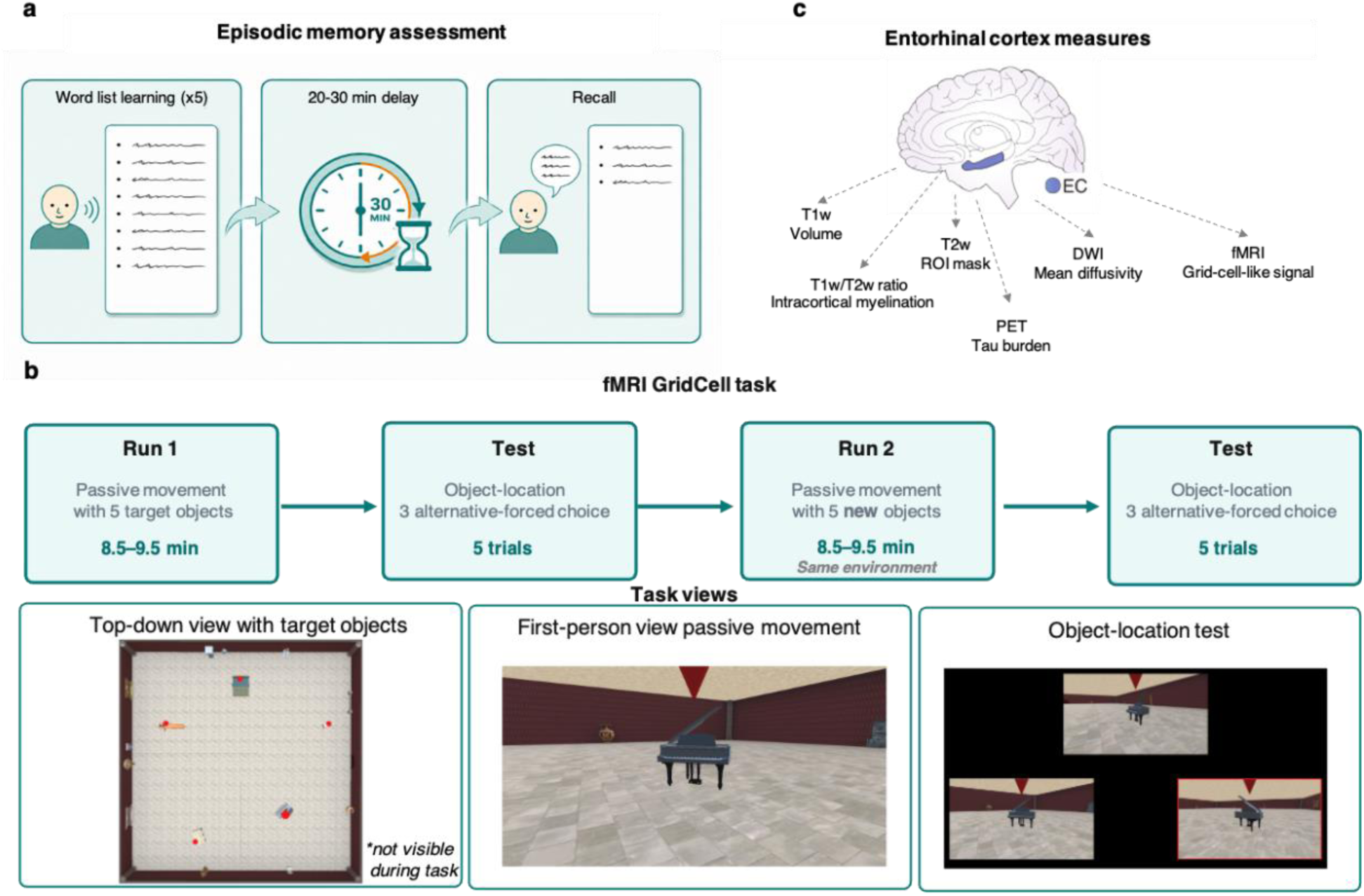
Study design and multimodal entorhinal cortex assessment. **a,** Verbal episodic memory was assessed using a word-list learning task (RAVLT). Participants learned a list of 15 words across five trials, this was followed by an intervening 15-word distractor list, immediate post-distractor recall, and a delayed recall test after 20-30 min which was used as an index of episodic memory **b**, During fMRI, participants completed a passive virtual-navigation task designed to elicit grid-cell-like representations. In each run, participants passively moved through a square virtual environment while encoding the locations of five target objects. Each navigation run was followed by a three-alternative forced-choice object-location test, where participants had to select the correct target location from three screenshots (two foils/one correct). Run two used a new set of objects within the same environment. Example screenshots show the first-person view during passive movement, the top-down environment with target objects, and the object-location test phase. **c,** Multimodal measures were extracted from the entorhinal cortex, including fMRI-derived grid-cell-like signal, PET-derived tau burden, and (micro-)structural MRI measures.

**Fig. 2.**
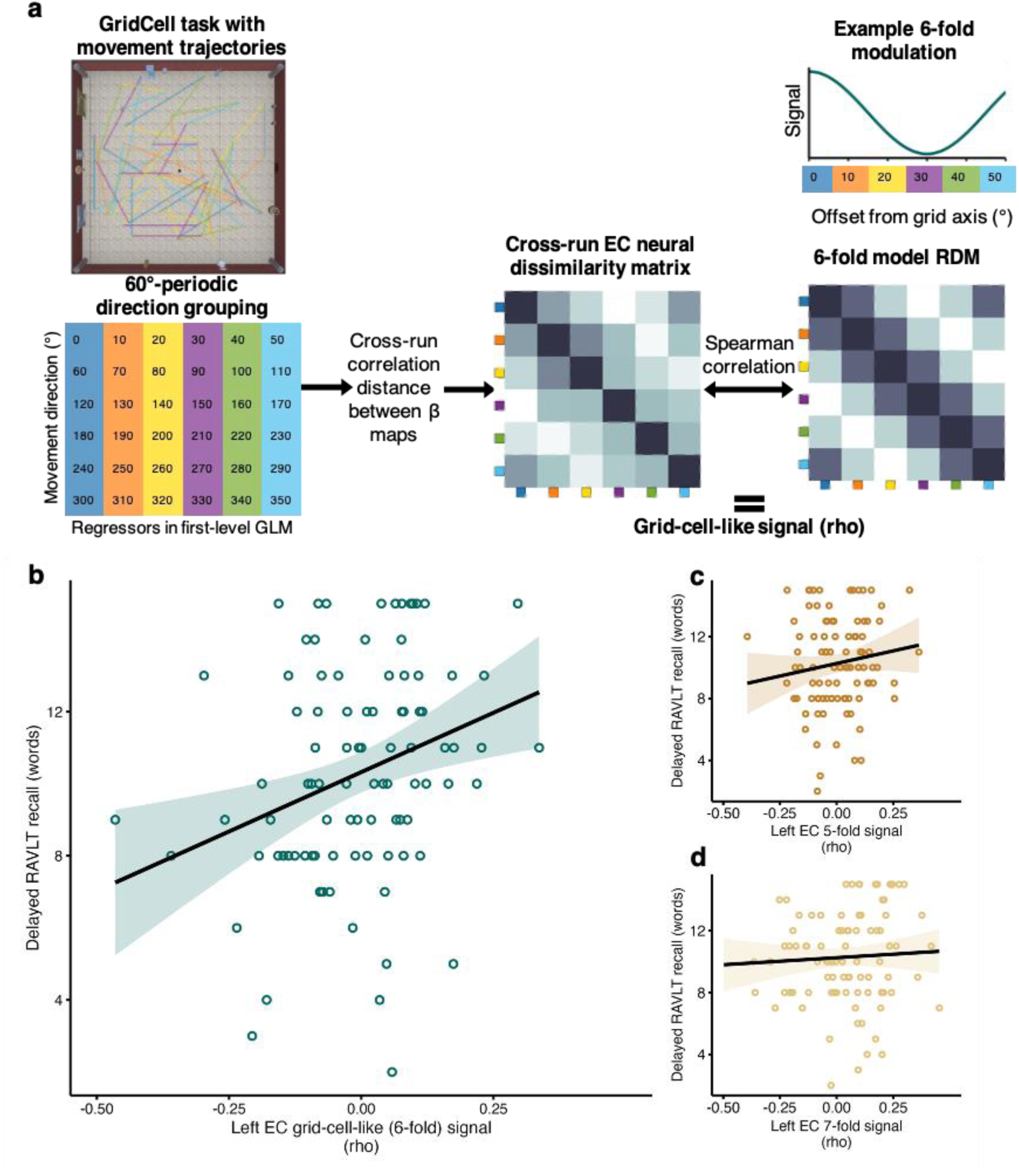
Left entorhinal grid-cell-like signal is associated with episodic memory performance. **a,** Schematic illustrating the multivariate representational similarity analysis used to estimate entorhinal grid-cell-like signal. Movement directions during virtual navigation were modelled as direction-specific regressors in the first-level GLM and grouped in 60° rotational space. The expected six-fold BOLD modulation is shown for an example grid orientation of 0°, with movement directions expressed relative to the grid axis. Cross-run correlation distances between direction-specific beta maps were used to construct entorhinal neural representational dissimilarity matrices (RDMs), which were compared with a six-fold model RDM using Spearman correlation. The resulting correlation coefficient provided an individual estimate of grid-cell-like signal. **b–d,** Scatter plots show the association between left entorhinal grid-cell-like signal and delayed word-list recall for the canonical six-fold signal (**b**) and control periodicities (**c,d**). Points show raw delayed-recall scores and raw grid-cell-like signal values. Regression lines show the model-estimated association adjusted for age, with age fixed at the sample mean.

We assessed the association between entorhinal grid-cell-like signal and episodic memory using Bayesian regression models with age as a covariate. This covariate was included because higher age was associated with lower delayed RAVLT recall in an age-only Bayesian regression model (unstandardized β = −0.12 words/year, 95% credible interval = −0.19 to −0.06, posterior probability of direction = 0.99). Bayes factors for the grid-cell-like signal analyses were obtained by comparing the full model, including entorhinal grid-cell-like signal and age, against an age-only null model. Analyses were conducted separately for left and right entorhinal cortex. Higher grid-cell-like signal in the left entorhinal cortex was associated with better delayed RAVLT recall after accounting for age (standardized β = 0.30, 95% credible interval = 0.11 to 0.49, posterior probability of direction = 0.99, Fig. 2b). Bayesian model comparison provided strong evidence that including left entorhinal grid-cell-like signal improved prediction of memory performance relative to an age-only model (BF₁₀ = 16.12). In contrast, grid-cell-like signal in the right entorhinal cortex was not positively associated with delayed RAVLT recall after accounting for age (standardized β = −0.10, 95% credible interval = −0.30 to 0.10, posterior probability of direction = 0.85; Supplementary Fig. S1). Bayesian model comparison did not support the inclusion of right entorhinal grid-cell-like signal over an age-only model (BF₁₀ = 0.43), indicating that the relationship between grid-cell-like activity and memory performance was specific to the left entorhinal cortex.

Since only grid-cell-like signal in the left entorhinal cortex showed evidence for an association with verbal episodic memory, subsequent analyses focused on the left entorhinal grid-cell signal as the memory-relevant grid-cell-like measure. To test whether the association with memory was specific to the canonical 6-fold grid-cell-like signal, we examined control periodicities in the left entorhinal cortex. Neither the 5-fold nor the 7-fold left entorhinal signal showed evidence for a comparable association with delayed RAVLT recall (Fig 2c-d). For the 5-fold signal, the effect was weak and uncertain (standardized β = 0.14, 95% credible interval = −0.07 to 0.34, posterior probability of direction = 0.91), with anecdotal evidence for including the predictor over the age-only model (BF₁₀ = 1.67). For the 7-fold signal, the effect was smaller and similarly uncertain (standardized β = 0.06, 95% credible interval = −0.14 to 0.26, posterior probability of direction = 0.72), with evidence favoring the age-only model over inclusion of the control periodicity (BF₁₀ = 0.31). Together, the memory association was only present for the 6-fold grid-cell-like signal, suggesting that preserved left entorhinal grid-cell-like representations are linked to better episodic memory performance in older adults.

### Entorhinal tau PET burden is associated with grid-cell–like coding

Having established that entorhinal grid-cell-like coding was related to episodic memory performance, we next tested whether entorhinal cortex tau accumulation was associated with reduced grid-cell-like signal. A subset of the participants (*n*=46) underwent dynamic tau PET imaging using the second-generation tracer [¹⁸F]PI-2620^31^ to quantify left entorhinal cortex tau burden (Fig 3a). Bayesian regression analysis showed a negative association between left entorhinal tau PET burden and local grid-cell-like signal after accounting for age (standardized β = −0.54, 95% credible interval = −0.81 to −0.28, posterior probability of direction = 0.99, Fig 3b). Bayesian model comparison provided strong evidence for including left entorhinal tau over the age-only model (BF₁₀ = 32.48). Importantly, this association was robust to the tau quantification approach: when left entorhinal tau burden was quantified using unadjusted distribution volume ratio (DVR) values, without white-matter correction, the negative association with grid-cell-like signal remained evident (β = −0.45, 95% CrI [−0.72, −0.17], BF₁₀ = 22.61).

**Fig. 3.**
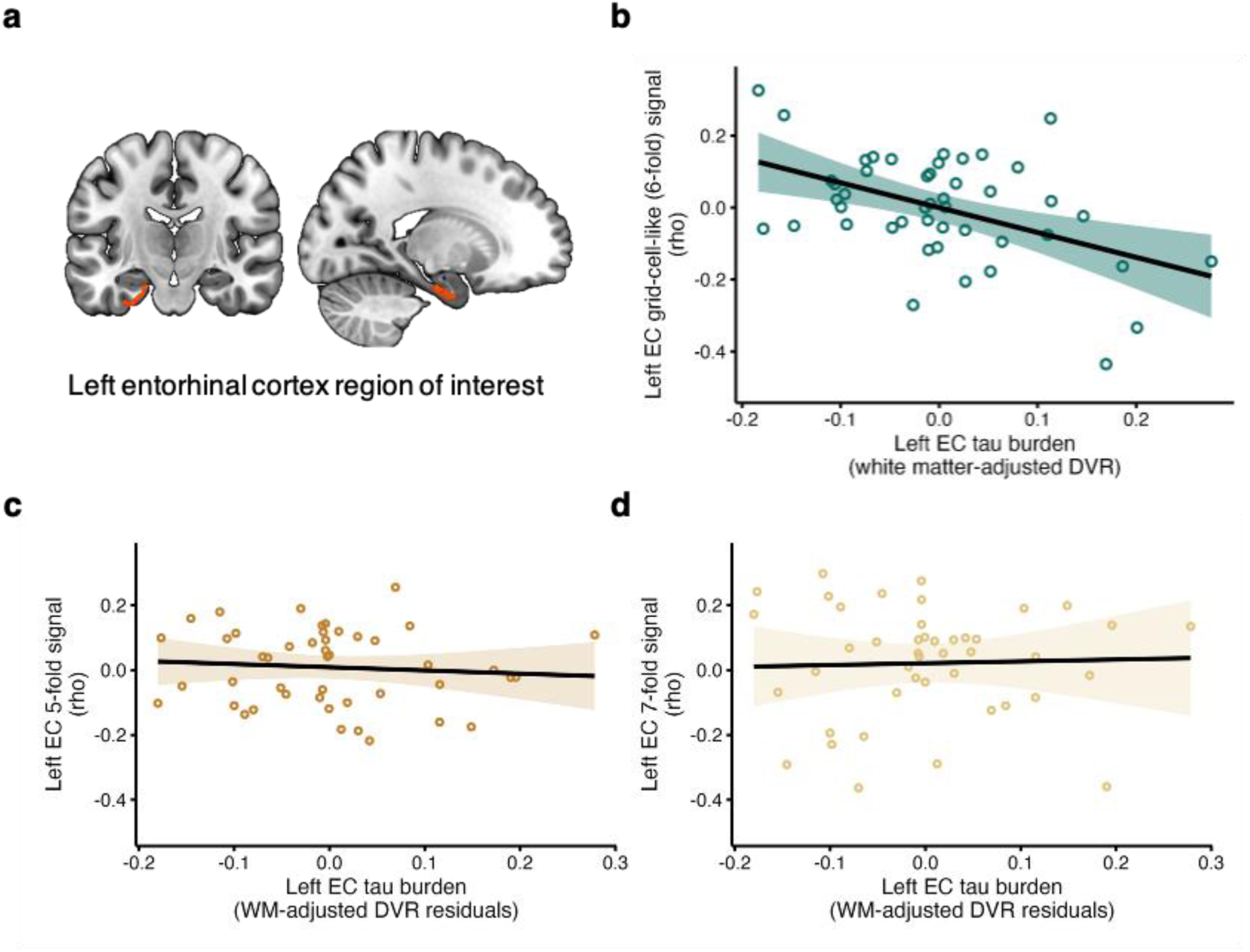
Entorhinal tau burden is associated with reduced grid-cell-like signal in the left entorhinal cortex. **a,** Left entorhinal cortex region of interest shown on coronal and sagittal anatomical slices. Left entorhinal cortex was segmented using OAP atlas^32^ and the Automatic Segmentation of Hippocampal Subfields software^33^. **b,** Higher left entorhinal tau burden, quantified as DVR adjusted for local white-matter binding, was associated with lower left entorhinal grid-cell-like signal at the canonical six-fold periodicity. **c, d,** Control analyses showed no corresponding associations between left entorhinal tau burden and non-canonical five-fold or seven-fold signals. Scatter plots show raw grid-cell-like signal values plotted against white-matter-adjusted left entorhinal DVR residuals. Regression lines show the model-estimated association adjusted for age, with age fixed at the sample mean. Shaded bands indicate 95% confidence intervals around the fitted regression line.

Further, to test whether this association was specific to grid-cell-like coding, we repeated the analysis for 5-fold and 7-fold control periodicities. Left entorhinal tau showed no comparable association with either the 5-fold signal (standardized β = −0.08, 95% credible interval = −0.38 to 0.22, BF₁₀ = 0.47, Fig 3c) or the 7-fold signal (standardized β = 0.03, 95% credible interval = −0.28 to 0.34, BF₁₀ = 0.42, Fig 3d). These findings indicate that the tau PET burden-related reduction in neural periodicity was specific to the 6-fold grid-cell-like signal.

### MRI-derived entorhinal tissue measures do not account for grid-cell-like signal

Given that tau accumulation may be accompanied by broader changes in regional structure^20^, and tissue integrity^34,35^ we next tested whether reduced grid-cell-like signal reflected a more general vulnerability of the left entorhinal cortex. Specifically, we examined whether individual differences in left entorhinal volume, diffusion-based microstructure, or T1w/T2w-derived intracortical tissue contrast were associated with left entorhinal six-fold grid-cell-like activity while controlling for age. These complementary analyses used all participants with available modality-specific data (volume: n = 100; T1w/T2w ratio: n = 100; mean diffusivity: n = 72). We found no evidence that left entorhinal volume, mean diffusivity, or T1w/T2w-derived intracortical tissue contrast were associated with left entorhinal six-fold grid-cell-like activity beyond age (all BF₁₀ < 1; volume: BF₁₀ = 0.34; mean diffusivity: BF₁₀ = 0.48; T1w/T2w ratio: BF₁₀ = 0.31; Supplementary Fig. S2). In addition, we also tested if chronological age was associated with left entorhinal six-fold grid-cell-like activity in the full fMRI sample. Age was not clearly associated with left entorhinal six-fold grid-cell-like signal (unstandardized β = 0.001, 95% credible interval = −0.002 to 0.005, posterior probability of direction = 0.75). Thus, the tau-related reduction in grid-cell-like coding was not mirrored by corresponding associations with chronological age or MRI-derived measures of entorhinal tissue integrity, suggesting that grid-cell-like signal was not simply a proxy for age, macroscopic atrophy, or microstructural alterations detectable with the present MRI measures.

### Exploratory evidence for an association between higher entorhinal tau and reduced representational stability

Rodent work in aged animals suggests that aging does not simply abolish spatially tuned cells but reduces the stability of their firing^16,17^. Early entorhinal tau pathology appears to affect a similar aspect of grid-cell coding, reducing grid periodicity and destabilizing grid fields^26–28^. Motivated by these findings, we conducted an exploratory analysis of temporal stability in our human fMRI data.

To this end, we estimated direction-related representational geometry separately for each run. For each run, we computed a neural representational dissimilarity matrix based on the multivoxel activity patterns associated with movement directions folded into 60° rotational space. Temporal stability was then quantified as the similarity between run 1 and run 2 neural RDMs (see Methods for more details). This measure therefore captured how consistently left entorhinal direction-related representational structure was expressed across runs, independently of its correspondence with the six-fold model RDM. Left entorhinal tau PET DVR showed a negative association with temporal stability in the predicted direction, although evidence for this effect was inconcslusive (β = −0.26, 95% CrI [−0.55, 0.02], pd = 0.97, BF₁₀ = 1.54, Supplementary Fig. S3). Thus, the observed pattern raises the possibility that tau accumulation may destabilize grid-cell-like representations over time.

### Medial temporal lobe tau relates to episodic memory through entorhinal grid-cell-like coding

Having shown that left entorhinal grid-cell-like signal was associated with episodic memory performance, and that left entorhinal tau burden predicted reduced grid-cell-like signal, we next tested whether grid-cell-like coding provides a functional pathway linking tau pathology to reduced episodic memory. Given that (i) the majority of previous work linking tau pathology to episodic memory in cognitively normal older adults has primarily focused on tau burden across the medial temporal lobe (MTL) ^21,36,37^, and (ii) episodic memory depends on distributed MTL circuitry^38,39^ rather than on the entorhinal cortex alone, we used MTL PET burden (mean DVR across entorhinal cortex, parahippocampal gyrus and fusiform gyrus, see Methods for details) as the predictor in our mediation analysis.

We conducted a Bayesian mediation analysis to test whether left entorhinal grid-cell-like signal provided an indirect pathway linking MTL tau burden to delayed word-list recall, while controlling for age (Fig. 4) Greater MTL tau PET burden was associated with reduced left entorhinal grid-cell-like signal (a-path: β = −0.44, 95% CrI [−0.71, −0.17], pd = 0.99), and stronger left entorhinal grid-cell-like signal was associated with better delayed recall when controlling for MTL and age (b-path: β = 0.30, 95% CrI [0.01, 0.59], pd = 0.98). The direct association between MTL and delayed recall was uncertain (β = −0.17, 95% CrI [−0.46, 0.12], pd = 0.88). Critically, the indirect effect through left entorhinal grid-cell-like signal was credible and in the expected negative direction (β = −0.13, 95% CrI [−0.31, −0.004], pd = 0.98). This negative indirect effect reflects the combination of a negative association between MTL tau burden and grid-cell-like signal, and a positive association between grid-cell-like signal and delayed recall. Thus, higher MTL tau burden was indirectly associated with lower delayed recall through reduced left entorhinal grid-cell-like signal. Sensitivity analyses showed a directionally similar but less certain mediation pattern when using unadjusted MTL DVR values, while the indirect effect was less clearly supported when tau burden was restricted to the local left entorhinal cortex (Supplementary Table 2). Together, these findings suggest that the MTL burden relates to episodic memory performance, at least in part, through the integrity of left entorhinal grid-cell-like coding.

**Fig. 4.**
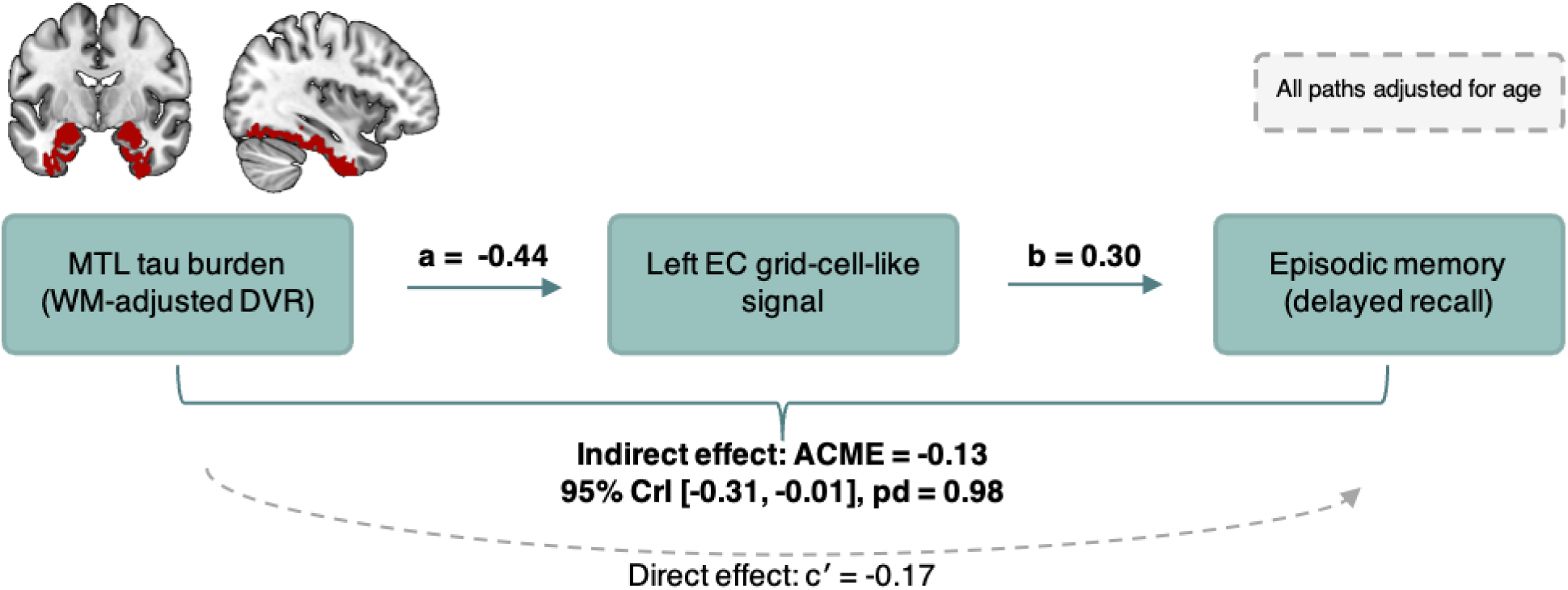
Mediation model linking medial temporal tau burden, entorhinal grid-cell-like signal and episodic memory. Bayesian mediation model testing whether left entorhinal cortex (EC) grid-cell-like signal mediates the association between medial temporal lobe (MTL) tau PET burden and episodic memory performance. The brain inset shows the MTL region used to quantify tracer binding. Age was included as a covariate in the mediator and outcome models. The model estimated both the indirect pathway through grid-cell-like signal and the direct association between MTL tau burden and episodic memory. pd, posterior probability of direction.

## Discussion

In this study, we tested whether entorhinal grid-cell-like representations provide a functional link between early tau pathology and episodic memory variability in normal aging. Combining virtual-reality fMRI, multivariate representational similarity analysis, tau PET imaging with the second-generation tau tracer [¹⁸F]PI-2620, and episodic memory assessment in cognitively normal older adults, we report three main findings. First, weaker left entorhinal grid-cell-like signal was associated with poorer episodic memory performance. Second, greater tau burden in the left entorhinal cortex was associated with reduced grid-cell-like signal. Third, Bayesian mediation analysis showed that medial temporal tau burden related to episodic memory performance indirectly through left entorhinal grid-cell-like signal. Together, these findings suggest that early tau accumulation in normal aging is associated with compromised entorhinal computation, and that disruption of grid-cell-like coding may constitute a functional pathway through which medial temporal tau pathology contributes to episodic memory differences in older adults.

Left entorhinal grid-cell-like signal was positively associated with episodic memory performance, such that individuals with weaker entorhinal six-fold coding exhibited lower episodic memory as measured using delayed verbal recall. Importantly, this relationship with episodic memory was specific to the canonical six-fold periodicity and was not observed for control periodicities, suggesting that the association reflected grid-cell-like coding rather than broader changes in entorhinal activity. The left-lateralized pattern observed here should be interpreted cautiously. Hemispheric lateralization of grid-cell-like coding was not a primary hypothesis, and human fMRI studies of grid-cell-like signals during navigation have not established a consistent lateralisation pattern^18,40,41^. The present left-sided association may instead reflect the verbal episodic-memory outcome used in this study, given broader evidence linking verbal memory to left-lateralized temporal-lobe systems^42^. This interpretation is also consistent with recent tau-PET findings suggesting similar material-dependent hemispheric associations between tau burden and memory performance, with verbal memory showing stronger disruption with left hemisphere tau burden^43^. Thus, the present findings should not be interpreted as evidence for a general left-hemispheric dominance of grid-cell-like coding but but rather as a possible material-dependent lateralization of the grid–verbal memory association.

A key question raised by these findings is why disruption of an entorhinal code measured during navigation would relate to delayed verbal recall. If grid-cell-like codes provide a spatial or relational scaffold for episodic memory^2^, then reduced grid-cell-like coding could weaken the organization of events within the contextual frameworks that support later retrieval. This account is consistent with object–location memory studies showing that forgetting can reflect swap errors^44^, in which objects are assigned to incorrect locations rather than forgotten entirely, pointing to a failure of spatial binding rather than item memory per se^45^. Thus, more broadly, disruption of grid-cell-like coding may render the spatial and contextual organization of experiences less reliable, weakening the scaffold through which events, locations, and contexts are bound into coherent episodic memories.

In the context of delayed word-list recall, this broader scaffolding function may be relevant because successful performance requires participants to bind words to a specific learning episode, maintain this context across a delay, distinguish the original list from competing material introduced by the interference list, and search memory in a structured way during retrieval. Free recall is not a random readout of stored items but is shaped by temporal and semantic organization: participants preferentially transition between items that were encoded close together in time and between items that are semantically related^46^ with these distances also reflected in hippocampal theta activity during recall^47^. This provides a direct link to grid-cell-like coding, because entorhinal grid-like representations have been observed not only during navigation, but also during the representation of semantic spaces, including word meaning^13^. On this view, reduced grid-cell-like coding could weaken the scaffold that helps organize items within their temporal, semantic, or contextual relationships, making the original learning episode less coherently structured or less efficiently searchable during delayed recall.

The second key finding of our study is that higher left entorhinal tau burden was associated with reduced grid-cell-like signal. The entorhinal cortex is among the first regions to show tau accumulation in aging and Alzheimer’s disease^25^, and previous work suggests that tau pathology in this region is related to functional brain changes even in cognitively healthy older adults, including altered, largely hyperactive^21,48,49^, medial temporal activity as well as disrupted network specificity^50,51^ and connectivity^20^. However, these studies have not established the neural computations through which entorhinal tau pathology may affect episodic memory. Our findings address this gap by showing that entorhinal tau burden in cognitively healthy older adults is associated with reduced grid-cell-like coding, a local entorhinal computation identified here as relevant to episodic memory performance. This is consistent with rodent work showing that tau pathology in the entorhinal cortex impairs grid-cell firing and spatial memory^26–28,52^. Importantly, our results extend this work to cognitively healthy older adults, in whom tau burden is relatively low and clinical impairment is absent. Thus, tau-related disruption of grid-cell-like coding may not only be a feature of overt disease, but may already be detectable during cognitively normal aging, when molecular pathology is emerging, but structural degeneration remains subtle.

The absence of corresponding associations with entorhinal volume, mean diffusivity, intracortical myelination is important in this context. These results suggest that the relationship between tau burden and grid-cell-like signal was not simply explained by detectable macroscopic atrophy, microstructural change, or intracortical myelin differences. Instead, tau pathology may impair grid-cell-like coding through synaptic^53–55^ and circuit-level mechanisms that are not necessarily captured by these MRI measures. This interpretation is consistent with evidence that aging affects hippocampal–entorhinal function through changes in synaptic plasticity, experience-dependent stability, and circuit dynamics, rather than through cell loss alone^56,57^. It is also consistent with mechanistic evidence that pathological tau can interfere directly with synaptic function^58^, including presynaptic vesicle mobility, release rate, and neurotransmission^59^, and with human evidence linking entorhinal tau deposition to lower hippocampal synaptic density in older individuals with normal cognition and early Alzheimer’s disease^60^. Such synaptic and circuit-level effects could degrade the precision, coordination, or reliability of entorhinal population codes, thereby reducing grid-cell-like signal without producing detectable alterations in regional volume, mean diffusivity, or intracortical myelination.

One possible circuit-level expression of tau-related dysfunction is reduced temporal stability of grid-cell-like representations. Rodent studies of aging suggest that spatially tuned neurons become less stable over time: hippocampal place-cell representations show reduced temporal stability in aged animals^17^, and entorhinal grid cells show weaker stabilization across experience^16^. Human fMRI work further suggests that age-related reductions in grid-cell-like signal are closely related to reduced temporal stability of grid orientation within the entorhinal cortex^16^. Consistent with this account, recent rodent work shows that tau pathology in the medial entorhinal cortex induces neuronal hyperactivity and increases representational instability, particularly when animals navigate familiar environments^28^. Rodent work further links entorhinal tau pathology to reduced grid-cell periodicity, altered medial entorhinal cortex interneuron firing, and enhanced theta power^26^. Motivated by this work, we conducted an exploratory analysis of temporal stability in the human fMRI data. Although the association between left entorhinal tau PET burden and temporal stability was in the predicted negative direction, evidence for this effect was inconclusive and should therefore be interpreted cautiously. Nevertheless, this result is consistent with the possibility that tau accumulation may compromise grid-cell-like coding by reducing the reliability with which entorhinal representational structure is expressed over time.

More broadly, if early tau accumulation contributes to synaptic dysfunction or excitatory–inhibitory imbalance, it could impair the temporal coordination required for stable grid-cell-like representations. Consistent with this idea, computational modelling of grid-cell-to-place-cell transformations suggests that synaptic disruption in entorhinal–hippocampal circuits is sufficient to produce abnormalities in place-cell function, including reduced across-session stability of place fields and reduced place map resolution^61^. In this sense, tau-related grid-cell disruption may represent one expression of broader early circuit stress within the MTL.

Finally, our mediation analysis suggested that medial temporal tau burden relates to episodic memory performance, at least in part, through differences in left entorhinal grid-cell-like coding. Notably, this indirect effect was evident for broader medial temporal tau PET burden but was less clearly supported when tau was restricted to the left entorhinal cortex. This pattern is consistent with the idea that episodic memory depends on distributed MTL computations rather than on the entorhinal cortex alone^38,39^. Grid-cell-like coding may provide a spatial or relational scaffold for memory representations, but this scaffold is unlikely to depend only on local entorhinal computations. Instead, grid-cell-like representations are shaped by interactions with wider medial temporal and parahippocampal circuits, including presubicular and parasubicular inputs conveying head-direction and orientation information^62^, hippocampal–subicular feedback carrying spatial-contextual information, and parahippocampal–perirhinal pathways that provide contextual and item-related inputs^62,63^. Tau accumulation across this broader network may therefore be particularly relevant for episodic memory because it could perturb both entorhinal grid-cell-like computations and the medial temporal inputs that help anchor these computations to objects, locations, and contexts.

Several limitations of our study warrant more detailed discussion. First, grid-cell-like signal was measured during navigation, whereas episodic memory was assessed using delayed word-list recall. The present findings therefore do not show that grid-like codes were directly engaged during word encoding or retrieval. Instead, they suggest that the integrity of an entorhinal computation measured during navigation is related to a broader ability to organize and retrieve episodic information. Moreover, although delayed word-list recall is a well-established and sensitive measure of verbal episodic memory^64^, it is less naturalistic than everyday episodic memory, which often requires the integration of item, spatial, temporal, and contextual information^65^. This contrasts with the navigation task used to estimate grid-cell-like signal, which captures a more spatially structured and multifaceted behaviour. Future studies should therefore test whether entorhinal grid-cell-like coding relates not only to verbal delayed recall, but also to the spatial and temporal components of episodic memory, including memory for where and when events occurred.

Second, the absence of corresponding associations with entorhinal volume, mean diffusivity, or intracortical myelination does not rule out more subtle structural alterations. Rather, particularly in cognitively normal aging, tau-related reductions in grid-cell-like coding may reflect subtle cellular or subvoxel structural alterations that are not captured by the MRI measures used here, such as changes in dendritic architecture, spine or synapse density, layer-specific neuronal integrity, mitochondrial function, or microvascular integrity. Future studies combining tau PET and grid-cell-like fMRI with CSF-derived synaptic integrity markers^66^, synaptic-density PET^67^, advanced diffusion imaging to quantify to soma and neurite architecture^68^ as well as the integrity of critical entorhinal input pathways, could help elucidate the cellular and circuit-level mechanisms through which tau accumulation disrupts grid-cell-like coding in normal aging.

In conclusion, the present findings identify entorhinal grid-cell-like coding as a functional mechanism through which early tau accumulation may contribute to episodic memory variability in healthy aging. By showing that higher tau burden in the left entorhinal cortex is associated with reduced local grid-cell-like signal, our results suggest that tau accumulation in one of the earliest tau-affected medial temporal regions is already linked to altered memory-relevant computation in cognitively healthy older adults. Importantly, this disruption was not explained by macroscopic or microstructural MRI measures, pointing to grid-cell-like coding as a sensitive functional readout of tau-related circuit vulnerability. Together with the mediation results, these findings suggest that age-related differences in episodic memory may arise, in part, from tau-related disruption of the entorhinal computations that organize spatial and contextual information. More broadly, this work positions entorhinal grid-cell-like coding as a mechanistically informative bridge between molecular pathology and cognitive decline, and as a potential functional target for future studies aimed at detecting and preserving memory function in aging.

## Methods

### Participants

Participants were recruited within the Collaborative Research Centre 1436 “Neural Resources of Cognition”, primarily from its previously described central healthy-aging cohort^69^, with additional recruitment through CRC 1436–associated studies. All participants provided informed consent, and the study was approved by the Ethics Committee of the University of Magdeburg. The present sample consisted of healthy, community-dwelling older adults who were screened for major neurological or psychiatric disease, major medical illness, and substance-related addiction. Because the study focused on normal cognitive aging, cognitive status was assessed using the Consortium to Establish a Registry for Alzheimer’s Disease Neuropsychological Battery Plus^70^ (CERAD-Plus), where available. Participants were excluded if they scored below 26 on the Mini-Mental State Examination, administered as part of the CERAD-Plus battery, or if their global CERAD-Plus performance fell below −1.5 SD relative to age-, sex-, and education-adjusted norms. Six participants did not complete the CERAD-Plus battery but had Montreal Cognitive Assessment data and scored above 23^71^, consistent with the recommended cut-off for normal cognition

A total of 102 cognitively unimpaired older adults underwent structural MRI. One participant was excluded because the entorhinal cortex could not be reliably segmented due to atypical anatomy, and one participant was excluded due to impaired vision problems that affected task performance. The final fMRI sample comprised 100 participants, including 50 female and 50 male participants, with a mean age of 76.3 years (SD = 7.64, range = 62–96 years). For the primary episodic memory analysis, 93 participants had both usable fMRI and verbal learning and memory test data. A subset of 72 participants had overlapping diffusion MRI data and was included in analyses of mean diffusivity. Tau PET data overlapping with usable grid-cell-like signal and episodic memory data were available for 46 participants and were used for the PET analyses. Demographic, cognitive, genetic, plasma biomarker, and PET characteristics of the sample and modality-specific subsamples are reported in Supplementary Table 1.

### Episodic memory assessment: verbal learning and memory test

Episodic memory was assessed using the German version of the Rey Auditory Verbal Learning Test (RAVLT)^33^. Participants were first presented with a list of 15 unrelated words, which were read aloud by the examiner at a constant rate. Immediately after the presentation, participants were asked to freely recall as many words as possible, in any order. This learning-and-recall procedure was repeated across five consecutive learning trials using the same word list. Participants were then presented with a second interference list of 15 new words and again asked to recall as many words as possible. Immediately after the interference trial, participants were asked to recall the original word list again without re-presentation. After a 20-30 minute delay, during which participants completed unrelated tasks, delayed free recall of the original word list was assessed. Following delayed free recall, participants completed a recognition trial in which they identified words from the original list among distractor items. The present analyses focused on delayed free recall as the primary episodic memory outcome.

### fMRI grid-cell task

Participants also performed a passive virtual navigation task designed to sample a broad range of travel directions for the assessment of grid-cell-like representations. In each run, participants were translated along one of nine predefined trajectory versions, which were pseudorandomized across participants. Each trajectory comprised translation periods, rotations, and brief stationary intervals, producing a random-walk-like path through the environment. Translation speed was held constant at 15 virtual m/s, rotations were performed at 20°/s, and translation segments varied in duration (mean = 3.45 s, SD = 0.75). Depending on the assigned trajectory, each run lasted between 8.5 and 9.5 min.

The virtual environment consisted of a square room (140 vm x 140 vm) with textured flooring and wall cues providing orientation information. Five everyday objects were distributed across the environment and marked by red cones to facilitate visibility. In the second run, the spatial layout remained identical, but the target objects were replaced. Participants were instructed to imagine actively moving through the environment and to encode the object locations. After each run, they completed a three-alternative forced-choice object-location test, selecting the screenshot that showed the correct location of each target object, with 5 test trials per run.

### MRI acquisition

Structural MRI and fMRI data were acquired on a 3T Siemens Prisma scanner using a 64-channel head coil. A whole-brain T1-weighted MPRAGE image was acquired with the following parameters: TR = 2,500 ms, TE = 4.37 ms, TI = 1,100 ms, flip angle = 7°, and 1.0 mm isotropic resolution. In addition, two T2-weighted images were acquired. A coronal T2-weighted turbo spin echo scan, oriented perpendicular to the hippocampal long axis, was acquired for entorhinal and MTL segmentation (TR = 6,000 ms, TE = 71 ms, 0.5 × 0.5 × 2.0 mm resolution). A separate high-resolution 3D T2-weighted SPACE ZOOMit slab acquisition focused on the inner ear was also acquired but was not used in the present analyses. Functional images were acquired using a two-echo 3D EPI sequence with the following parameters: TR = 2,200 ms, TE1 = 14.2 ms, TE2 = 39.2 ms, flip angle = 15°, 2.0 mm isotropic resolution, 80 slices, and CAIPIRINHA acceleration of 3 × 2. Two task runs were acquired, each comprising 276 volumes.

Diffusion MRI data, available for a subset of 72 participants, were acquired in a separate session on a Siemens Skyra scanner using a 32-channel head coil (full details described in Behrenbruch^72^). Diffusion-weighted images were acquired using a single-shot echo-planar imaging sequence with the following parameters: TR = 4,010 ms, TE = 72.4 ms, 2.0 mm isotropic resolution, field of view = 240 × 240 × 144 mm³, two-fold parallel acceleration, and partial Fourier factor = 7/8. The acquisition included 80 volumes with b-values of 0, 700, and 1,000 s/mm², comprising 10 b = 0 volumes and 30 diffusion directions for each non-zero b-value.

### Entorhinal cortex segmentation

Segmentation of the entorhinal cortex was carried out with the Automatic Segmentation of Hippocampal Subfields (ASHS) pipeline^33^, using the ASHS-OAP atlas^32^, an atlas developed from older adults for automated segmentation of MTL cortical regions and hippocampal subfields. This procedure yielded separate masks for the anterolateral and posteromedial entorhinal cortex in each hemisphere. An experienced rater (PB) then quality-controlled the resulting masks against the manual segmentation protocol of Olsen et al^73^. One participant was excluded because the entorhinal cortex could not be reliably segmented due to anatomical variability. All remaining entorhinal masks passed visual quality control.

For the present analyses, anterolateral and posteromedial entorhinal masks were combined within each hemisphere to create total left and right entorhinal cortex masks. We used the total entorhinal cortex masks as the spatial resolution of the PET data made subregional entorhinal PET analyses less appropriate. Applying the same participant-specific total entorhinal masks across modalities ensured that fMRI, PET, structural MRI, T1w/T2w ratio, and diffusion-derived measures were extracted from anatomically corresponding entorhinal cortex regions.

### fMRI preprocessing and first-level modelling

Functional MRI preprocessing was performed using SPM12^74^ in MATLAB. Motion parameters were estimated from the first echo and the resulting transformations were applied to the second echo, after which all volumes were resliced. The two echoes were combined using voxel-wise weights derived from a brief resting-state scan acquired with the same sequence parameters, comprising 80 volumes. For each echo, temporal signal-to-noise ratio maps were computed and used to generate voxel-wise echo-combination weights. These weights were applied to each volume, yielding one optimally combined functional time series per run. The combined images were spatially smoothed with a 5 mm full-width-at-half-maximum Gaussian kernel.

Physiological noise regressors were derived using an anatomical CompCor approach^75^. The T1-weighted image was coregistered to the mean EPI image and segmented into gray matter, white matter, and cerebrospinal fluid using SPM’s unified segmentation procedure. The first five principal components extracted from white matter and cerebrospinal fluid masks were retained as nuisance regressors. The T2-weighted image and entorhinal cortex masks were coregistered to the participant’s EPI data to enable native-space ROI analyses.

First-level general linear models were estimated for each participant and included direction-specific regressors defined from periods of forward translation through the virtual environment. For each translation segment, the instantaneous movement direction was expressed as an angular heading and folded into 60° rotational space, reflecting the six-fold rotational symmetry characteristic of grid-cell firing. This yielded six direction regressors corresponding to 0°, 10°, 20°, 30°, 40°, and 50° in 60° rotational space, with each regressor pooling directions separated by multiples of 60°. Regressors were modelled using the onset and duration of the corresponding translation periods and convolved with the canonical hemodynamic response function. Each model additionally included the six realignment parameters and five anatomical CompCor components as nuisance regressors. Model estimation yielded one beta image for each direction regressor in each run, which formed the basis for the representational similarity analyses of grid-cell-like signal and temporal stability.

### Multivariate grid-cell-like analysis

Grid-cell–like signal was assessed using multivariate representational similarity analysis^76^. For each participant and ROI, voxel-wise beta estimates from the six direction regressors were extracted separately for run 1 and run 2. We then computed a cross-run neural representational dissimilarity matrix (RDM) by comparing each direction-specific multivoxel pattern from run 1 with each direction-specific pattern from run 2 using correlation distance. This yielded a 6 × 6 neural RDM for each ROI, in which each cell reflected the dissimilarity between two movement-direction bins estimated from independent runs. Computing dissimilarities across runs reduced the influence of within-run noise structure and temporal autocorrelation on the neural RDM.

To test whether this neural representational structure followed the six-fold symmetry expected for grid-cell-like coding, we constructed a model RDM based on the angular difference between movement-direction bins. Angular differences were transformed using a cosine function with six-fold periodicity, such that directions separated by multiples of 60° were predicted to have more similar activity patterns: (RDM₍model₎ = 1 − cos(6Δ*θ*)). The neural and model RDMs were vectorized and compared using Spearman rank correlation. The resulting correlation coefficient (rho) served as the participant-specific estimate of grid-cell-like signal strength for each region of interest.

### Exploratory temporal stability of entorhinal representational geometry analysis

To examine whether tau PET burden was associated with the temporal stability of entorhinal coding, we conducted an exploratory analysis in the left entorhinal cortex, where the main tau-related reduction in grid-cell-like signal was observed. Using the same first-level GLM outputs as the grid-cell-like analysis, we computed separate 6 × 6 neural RDMs for run 1 and run 2 based on pairwise correlation distances between the six direction-specific multivoxel beta patterns.

The off-diagonal elements of the two run-specific RDMs were then vectorized and correlated across runs. The resulting correlation coefficient indexed how consistently direction-related representational geometry was expressed across the two runs. Although this measure was computed in 60° rotational space, it was model-free and did not require correspondence with the six-fold grid-cell-like model RDM. Thus, the analysis tested the temporal reliability of entorhinal direction-related representational structure independently of grid-cell-like signal strength.

### Tau PET acquisition, processing, and white-matter adjustment

Tau PET acquisition and preprocessing followed procedures described previously for the healthy-aging cohort from which the present PET subsample was partly drawn^23^. Briefly, dynamic [¹⁸F]PI-2620 PET data were acquired simultaneously with MRI on a Siemens Biograph MR PET-MRI scanner. Participants received an intravenous bolus injection of [¹⁸F]PI-2620 (mean = 183.29 MBq, SD = 1.84, range = 175.9– 184.9), followed by 60 min of dynamic PET acquisition. Data were reconstructed into 34 frames and corrected for attenuation using MR-derived dual-echo ultra-short echo time segmentation, with the vendor Brain-HiRes protocol used when UTE segmentation failed. The dynamic PET series was motion-corrected and coregistered to the same-session T1-weighted image (MPRAGE; TR = 2.5 s, TE = 4.37 ms, TI = 1.1 s, flip angle = 7°, 1 mm isotropic).

Parametric BP_ND_ maps were estimated using the Multilinear Reference Tissue Model 2 (MRTM2^77^), as implemented in QModeling (v3; MATLAB R2024b), with inferior cerebellar grey matter as the reference tissue BP_ND_ values were converted to DVR by adding 1.

For the entorhinal cortex analyses, participant-specific entorhinal cortex masks were registered to the T1-weighted image acquired during the MR-PET session, allowing PET signal to be extracted from the same anatomically defined entorhinal cortex regions used in the fMRI analyses. To control for local white-matter binding of [¹⁸F]PI-2620, we created a subject-specific entorhinal cortex-adjacent white-matter mask by dilating the left total entorhinal cortex mask by 9 voxels, intersecting it with the subject-specific white-matter segmentation, and eroding the resulting mask by 2 voxels to reduce partial-volume contamination from neighbouring grey matter. Mean BP_ND_ from this entorhinal-adjacent white-matter mask was converted to DVR and used as a participant-specific estimate of local white-matter binding (see also Maass^21^). White-matter-adjusted left entorhinal cortex tau burden was defined as the residual from a linear regression in which left entorhinal cortex DVR was predicted by entorhinal cortex-adjacent white-matter DVR.

For the mediation analysis, PET signal was additionally extracted from a broader bilateral MTL ROI derived from FreeSurfer 7.1 segmentation of the same-session T1-weighted image using the Desikan– Killiany atlas^78^. This ROI comprised bilateral entorhinal cortex, parahippocampal cortex, and fusiform gyrus. The hippocampus was excluded to minimize potential contamination of its PET signal by spill-in from off-target tracer binding in the adjacent choroid plexus, a recognized concern in tau-PET quantification^23,79^. For this MTL ROI, white-matter adjustment was based on global FreeSurfer-derived whole-brain white-matter DVR, as described previously^21^. White-matter-adjusted MTL tau burden was defined as the residual from a linear regression in which MTL DVR was predicted by global white-matter DVR. These white-matter-adjusted DVR measures were used as the primary PET metrics. Sensitivity analyses were performed using the corresponding unadjusted DVR values for the left entorhinal cortex and MTL ROIs.

### Structural and diffusion MRI control measures

To extract left entorhinal cortex volume, raw volumes were obtained from the masks generated by the ASHS “UseGray” algorithm, which yields gray-matter-constrained volumetric estimates. To account for individual differences in head size, these raw volumes were corrected for total intracranial volume. Corrected values were taken as the residuals of a linear regression in which raw volume was the dependent variable and intracranial volume the predictor. The residualised volumes thus index each region’s volume independently of head size and served as the input for all downstream analyses.

Intracortical myelination was measured by the voxelwise T1w/T2w intensity ratio, which is commonly interpreted as being sensitive to intracortical myelin content^33^. The T1w image was first segmented using SPM12 unified segmentation, producing tissue classes and a bias-field correction. The T2w image was rigidly coregistered to the T1w using normalized mutual information as the cost function, resliced with fourth-order B-spline interpolation, and likewise bias-corrected. Only these coregistered, bias-corrected images were carried forward. Each image was divided by the median signal within its white-matter mask. Voxelwise T1w/T2w ratio maps were then computed within the gray-matter mask, with higher values reflecting greater myelin content. Finally, the entorhinal cortex masks were resliced to the processed T1w using nearest-neighbour interpolation, and the mean ratio within each subregion was extracted.

The preprocessing of dMRI data was performed using established protocols and open-source software from MRtrix3 and FSL consisting of denoising, correcting for susceptibility-induced geometric distortions (using opposite phase-encoding b0 images), and correcting for motion and Eddy-current artifacts. Bias field correction was applied using the ANTs algorithm via MRtrix3’s *dwibiascorrect*, and brain masks were generated with FSL’s *BET*. Eddy QC metrics were also generated using *eddy_quad*.

Following preprocessing, diffusion tensor reconstruction was performed using DIPY through *dipy.reconst.dti.TensorModel* generating mean diffusivity (MD) maps. The entorhinal masks were moved into diffusion space. Specifically, the T2w image and the entorhinal cortex masks were first registered to a mean b0 volume, and the resulting transform was applied to the masks and resampled onto the diffusion grid. MD was then computed as the partial-volume-weighted mean over all voxels with nonzero mask weight.

### Statistical information

All statistical analyses were conducted in RStudio. Bayesian linear regression models were fitted using the brms package with a Gaussian likelihood to test whether: (i) grid-cell-like signal predicted delayed RAVLT recall; (ii) regional tau burden predicted grid-cell-like signal; (iii) left entorhinal structural or microstructural measures predicted grid-cell-like signal; and (iv), in an exploratory analysis, left entorhinal tau burden predicted the temporal stability of left entorhinal representational geometry. The same framework was used in control analyses testing whether associations with delayed RAVLT recall and tau burden were specific to the canonical six-fold signal or were also present for non-grid control periodicities.

All models included age as a covariate and were fitted using complete cases for the variables included in each model. Standardized beta coefficients were obtained from models in which the outcome, predictor, and covariate were z-standardized before model estimation. For each Bayesian regression, we report the posterior mean standardized beta coefficient, 95% credible interval, and posterior probability of direction, defined as the posterior probability that the effect was positive or negative, whichever was larger. Model convergence was assessed using R^ and effective sample size diagnostics. Bayes factors were computed using the BayesFactor package by comparing each full model, including the predictor of interest and age, against an age-only null model. BF₁₀ therefore quantifies whether the predictor of interest improved model evidence beyond age.

Bayesian mediation analyses were performed to test whether left entorhinal grid-cell-like signal mediated the relationship between MTL tau burden and episodic memory performance. The primary mediation model used white-matter-adjusted bilateral MTL tau burden as the predictor, left entorhinal grid-cell-like signal as the mediator, and delayed word-list recall as the outcome. Left entorhinal tau burden was examined in an additional supplementary mediation analysis. For each mediation model, all variables were z-scored before analysis. We fitted two Bayesian Gaussian regression models using *brms*: a mediator model predicting left entorhinal grid-cell-like signal from MTL tau burden and age, and an outcome model predicting delayed word-list recall from left entorhinal grid-cell-like signal, MTL tau burden, and age. Weakly informative priors were used for all regression coefficients, β ∼ Normal(0, 1), with Student-t(3, 0, 2.5) priors for intercepts and Exponential(1) priors for residual standard deviations. Models were fitted with four chains, 4,000 iterations per chain, and 1,000 warm-up iterations, using an adapt_delta value of 0.95.

Posterior samples were extracted from both models to estimate the mediation paths. The **a** path was defined as the posterior distribution of the association between MTL tau burden and left entorhinal grid-cell-like signal. The **b** path was defined as the posterior distribution of the association between left entorhinal grid-cell-like signal and delayed word-list recall, controlling for MTL tau burden and age. The indirect effect was computed for each posterior draw as the product of the corresponding a and b path samples. The direct effect was defined as the posterior distribution of the MTL tau coefficient in the outcome model, and the total effect was computed as the sum of the direct and indirect effects. The same procedure was used for supplementary mediation analyses using local left entorhinal tau burden and for sensitivity analyses using unadjusted MTL DVR values. For each effect, we report the posterior mean, median, 95% credible interval, and posterior probability of direction.

## Data Availability

The datasets generated and/or analysed during the current study are not publicly available due to the inclusion of sensitive participant information and privacy concerns but are available from the corresponding author upon request.

## Code Availability

The code used for the multivariate analysis of the fMRI data, including the generation of the grid-cell-like signal and the temporal-stability metric, is publicly available on Zenodo at https://doi.org/10.5281/zenodo.21915222. Additional code used for statistical analyses is available from the corresponding authors upon request

## Acknowledgements

This work was supported by the VELUX Stiftung (Project No. 1809) and the German Research Foundation (Deutsche Forschungsgemeinschaft, DFG) under Project IDs 425899996 (CRC 1436) and 362321501 (RTG 2413). We thank Jiayu Chen for valuable contributions to the discussion and interpretation of the results. We also gratefully acknowledge Malika Schaumburg, Julia-Isabel Schinke, Berkant Bay, Felix Zeller who contributed to project administration and data collection.

## References

1. Rönnlund, M., Nyberg, L., Bäckman, L. & Nilsson, L.-G. Stability, Growth, and Decline in Adult Life Span Development of Declarative Memory: Cross-Sectional and Longitudinal Data From a Population-Based Study. Psychol. Aging 20, 3–18 (2005).

2. Sugar, J. & Moser, M.-B. Episodic memory: Neuronal codes for what, where, and when. Hippocampus 29, 1190–1205 (2019).

3. Moser, M.-B., Rowland, D. C. & Moser, E. I. Place Cells, Grid Cells, and Memory. Cold Spring Harb. Perspect. Biol. 7, a021808 (2015).

4. Solomon, E. A. et al. Dynamic Theta Networks in the Human Medial Temporal Lobe Support Episodic Memory. Curr. Biol. 29, 1100–1111.e4 (2019).

5. Dickerson, B. C. & Eichenbaum, H. The Episodic Memory System: Neurocircuitry and Disorders. Neuropsychopharmacology 35, 86–104 (2010).

6. Carr, V. A. et al. Individual differences in associative memory among older adults explained by hippocampal subfield structure and function. Proc. Natl. Acad. Sci. 114, 12075–12080 (2017).

7. Yassa, M. A., Mattfeld, A. T., Stark, S. M. & Stark, C. E. L. Age-related memory deficits linked to circuit-specific disruptions in the hippocampus. Proc. Natl. Acad. Sci. U. S. A. 108, 8873–8878 (2011).

8. Trelle, A. N., Henson, R. N. & Simons, J. S. Neural evidence for age-related differences in representational quality and strategic retrieval processes. Neurobiol. Aging 84, 50–60 (2019).

9. Reagh, Z. M. et al. Functional Imbalance of Anterolateral Entorhinal Cortex and Hippocampal Dentate/CA3 Underlies Age-Related Object Pattern Separation Deficits. Neuron 97, 1187–1198.e4 (2018).

10. Hunsaker, M. R., Chen, V., Tran, G. T. & Kesner, R. P. The medial and lateral entorhinal cortex both contribute to contextual and item recognition memory: A test of the binding ofitems and context model. Hippocampus 23, 380–391 (2013).

11. Object and Place Memory in the Macaque Entorhinal Cortex | Journal of Neurophysiology | American Physiological Society. https://journals.physiology.org/doi/full/10.1152/jn.1997.78.2.1062.

12. Constantinescu, A. O., O’Reilly, J. X. & Behrens, T. E. J. Organizing conceptual knowledge in humans with a gridlike code. Science 352, 1464–1468 (2016).

13. Viganò, S., Rubino, V., Soccio, A. D., Buiatti, M. & Piazza, M. Grid-like and distance codes for representing word meaning in the human brain. NeuroImage 232, 117876 (2021).

14. Nitsch, A., Garvert, M. M., Bellmund, J. L. S., Schuck, N. W. & Doeller, C. F. Grid-like entorhinal representation of an abstract value space during prospective decision making. Nat. Commun. 15, 1198 (2024).

15. Huber, D. E. A memory model of rodent spatial navigation in which place cells are memories arranged in a grid and grid cells are non-spatial. eLife 13, RP95733 (2025).

16. Herber, C. S., Pratt, K. J. B., Shea, J. M., Villeda, S. A. & Giocomo, L. M. Spatial coding dysfunction and network instability in the aging medial entorhinal cortex. Nat. Commun. 16, 8770 (2025).

17. Barnes, C. A., Suster, M. S., Shen, J. & McNaughton, B. L. Multistability of cognitive maps in the hippocampus of old rats. Nature 388, 272–275 (1997).

18. Stangl, M. et al. Compromised Grid-Cell-like Representations in Old Age as a Key Mechanism to Explain Age-Related Navigational Deficits. Curr. Biol. 28, 1108–1115.e6 (2018).

19. Ossenkoppele, R. et al. Tau PET positivity in individuals with and without cognitive impairment varies with age, amyloid-β status, APOE genotype and sex. Nat. Neurosci. 28, 1610–1621 (2025).

20. Berron, D. et al. Early stages of tau pathology and its associations with functional connectivity, atrophy and memory. Brain 144, 2771–2783 (2021).

21. Marks, S. M., Lockhart, S. N., Baker, S. L. & Jagust, W. J. Tau and â-Amyloid Are Associated with Medial Temporal Lobe Structure, Function, and Memory Encoding in Normal Aging. J. Neurosci. Off. J. Soc. Neurosci. 37, 3192–3201 (2017).

22. Maass, A. et al. Entorhinal Tau Pathology, Episodic Memory Decline, and Neurodegeneration in Aging. J. Neurosci. Off. J. Soc. Neurosci. 38, 530–543 (2018).

23. Maass, A. et al. Associations of [18F]PI-2620 Binding with Memory and Phosphorylated Tau 217 in Cognitively Unimpaired Older Adults. J. Nucl. Med. 10.2967/jnumed.125.271927 (2026) doi:10.2967/jnumed.125.271927.

24. Cho, H. et al. In vivo cortical spreading pattern of tau and amyloid in the Alzheimer disease spectrum. Ann. Neurol. 80, 247–258 (2016).

25. Braak, H. & Braak, E. Neuropathological stageing of Alzheimer-related changes. Acta Neuropathol. (Berl.) 82, 239–259 (1991).

26. Fu, H. et al. Tau Pathology Induces Excitatory Neuron Loss, Grid Cell Dysfunction, and Spatial Memory Deficits Reminiscent of Early Alzheimer’s Disease. Neuron 93, 533–541.e5 (2017).

27. Ying, J. et al. Disruption of the grid cell network in a mouse model of early Alzheimer’s disease. Nat. Commun. 13, 886 (2022).

28. Rodriguez, G. A. et al. Impaired spatial coding and neuronal hyperactivity in the medial entorhinal cortex of aged APP knock-in mice. Cell Rep. 45, 117505 (2026).

29. Abrous, D. N. et al. Hallmarks of healthy cognitive aging: Inter-individual differences in aging trajectories. Ageing Res. Rev. 119, 103102 (2026).

30. Helmstaedter, C. & Durwen, H. F. VLMT: Verbaler Lern-und Merkfähigkeitstest: Ein praktikables und differenziertes Instrumentarium zur Prüfung der verbalen Gedächtnisleistungen. [VLMT: A useful tool to assess and differentiate verbal memory performance.]. Schweiz. Arch. Für Neurol. Neurochir. Psychiatr. 141, 21–30 (1990).

31. Kroth, H. et al. Discovery and preclinical characterization of [18F]PI-2620, a next-generation tau PET tracer for the assessment of tau pathology in Alzheimer’s disease and other tauopathies. Eur. J. Nucl. Med. Mol. Imaging 46, 2178–2189 (2019).

32. Mazloum-Farzaghi, N., Barense, M. D., Ryan, J. D., Stark, C. E. L. & Olsen, R. K. The Effect of Segmentation Method on Medial Temporal Lobe Subregion Volumes in Aging. Hum. Brain Mapp. 45, e70054 (2024).

33. Yushkevich, P. A. et al. Quantitative comparison of 21 protocols for labeling hippocampal subfields and parahippocampal subregions in in vivo MRI: Towards a harmonized segmentation protocol. NeuroImage 111, 526–541 (2015).

34. Gagliardi, G. et al. Cortical microstructural changes predict tau accumulation and episodic memory decline in older adults harboring amyloid. Commun. Med. 3, 106 (2023).

35. Reijner, N. et al. T1-weighted/T2-weighted ratio reflects microstructural changes in Alzheimer’s disease. Alzheimers Res. Ther. 18, 175 (2026).

36. Simon, S. S. et al. In vivo tau is associated with change in memory and processing speed, but not reasoning, in cognitively unimpaired older adults. Neurobiol. Aging 133, 28–38 (2024).

37. Pelgrim, T. A. D., Beran, M., Twait, E. L., Geerlings, M. I. & Vonk, J. M. J. Cross-sectional associations of tau protein biomarkers with semantic and episodic memory in older adults without dementia: A systematic review and meta-analysis. Ageing Res. Rev. 71, 101449 (2021).

38. Squire, L. R. & Zola-Morgan, S. The Medial Temporal Lobe Memory System. Science 253, 1380–1386 (1991).

39. Ranganath, C. A unified framework for the functional organization of the medial temporal lobes and the phenomenology of episodic memory. Hippocampus 20, 1263–1290 (2010).

40. Horner, A. J., Bisby, J. A., Zotow, E., Bush, D. & Burgess, N. Grid-like Processing of Imagined Navigation. Curr. Biol. 26, 842–847 (2016).

41. Doeller, C. F., Barry, C. & Burgess, N. Evidence for grid cells in a human memory network. Nature 463, 657–661 (2010).

42. Lee, T. M. C., Yip, J. T. H. & Jones-Gotman, M. Memory Deficits after Resection from Left or Right Anterior Temporal Lobe in Humans: A Meta-Analytic Review. Epilepsia 43, 283–291 (2002).

43. Arias, J. F. et al. Hemispheric lateralization of Tau associations with visual and verbal memory in Alzheimer’s disease. Eur. J. Nucl. Med. Mol. Imaging 10.1007/s00259-026-07898-z (2026) doi:10.1007/s00259-026-07898-z.

44. Pertzov, Y., Dong, M. Y., Peich, M.-C. & Husain, M. Forgetting What Was Where: The Fragility of Object-Location Binding. PLOS ONE 7, e48214 (2012).

45. Segen, V., Avraamides, M. N., Slattery, T. J. & Wiener, J. M. Age-related differences in visual encoding and response strategies contribute to spatial memory deficits. Mem. Cognit. 49, 249–264 (2021).

46. Howard, M. W. & Kahana, M. J. When Does Semantic Similarity Help Episodic Retrieval? J. Mem. Lang. 46, 85–98 (2002).

47. Solomon, E. A., Lega, B. C., Sperling, M. R. & Kahana, M. J. Hippocampal theta codes for distances in semantic and temporal spaces. Proc. Natl. Acad. Sci. 116, 24343–24352 (2019).

48. Berron, D. et al. Higher CSF Tau Levels Are Related to Hippocampal Hyperactivity and Object Mnemonic Discrimination in Older Adults. J. Neurosci. 39, 8788–8797 (2019).

49. Huijbers, W. et al. Tau Accumulation in Clinically Normal Older Adults Is Associated with Hippocampal Hyperactivity. J. Neurosci. 39, 548–556 (2019).

50. Harrison, T. M. et al. Tau deposition is associated with functional isolation of the hippocampus in aging. Nat. Commun. 10, 4900 (2019).

51. Maass, A. et al. Alzheimer’s pathology targets distinct memory networks in the ageing brain. Brain 142, 2492–2509 (2019).

52. Ridler, T., Witton, J., Phillips, K. G., Randall, A. D. & Brown, J. T. Impaired speed encoding and grid cell periodicity in a mouse model of tauopathy. eLife 9, (2020).

53. Daniel Estrella, L., et al. Tau association with synaptic mitochondria coincides with energetic dysfunction and excitatory synapse loss in the P301S tauopathy mouse model. Neurobiol. Aging 147, 163–175 (2025).

54. Yoshiyama, Y. et al. Synapse Loss and Microglial Activation Precede Tangles in a P301S Tauopathy Mouse Model. Neuron 53, 337–351 (2007).

55. Hoover, B. R. et al. Tau Mislocalization to Dendritic Spines Mediates Synaptic Dysfunction Independently of Neurodegeneration. Neuron 68, 1067–1081 (2010).

56. Barnes, C. A., Srivathsa, S., Hill, P. F., Srokova, S. & Ekstrom, A. D. How Does Aging Impact the Structure and Function of Parallel Navigation Circuits across Mammals? in Challenges in Navigation Research: Mapping New Directions (eds Newcombe, N. S. & Cheng, K.) 321–378 (Springer Nature Switzerland, Cham, 2026). doi:10.1007/978-3-032-20563-6_13.

57. Shen, J., Barnes, C. A., McNaughton, B. L., Skaggs, W. E. & Weaver, K. L. The Effect of Aging on Experience-Dependent Plasticity of Hippocampal Place Cells. J. Neurosci. 17, 6769–6782 (1997).

58. Wu, M. et al. The role of pathological tau in synaptic dysfunction in Alzheimer’s diseases. Transl. Neurodegener. 10, 45 (2021).

59. Zhou, L. et al. Tau association with synaptic vesicles causes presynaptic dysfunction. Nat. Commun. 8, 15295 (2017).

60. Mecca, A. P. et al. Association of entorhinal cortical tau deposition and hippocampal synaptic density in older individuals with normal cognition and early Alzheimer’s disease. Neurobiol. Aging 111, 44–53 (2022).

61. Ness, N. & Schultz, S. R. A computational grid-to-place-cell transformation model indicates a synaptic driver of place cell impairment in early-stage Alzheimer’s Disease. PLOS Comput. Biol. 17, e1009115 (2021).

62. Canto, C. B., Koganezawa, N., Beed, P., Moser, E. I. & Witter, M. P. All Layers of Medial Entorhinal Cortex Receive Presubicular and Parasubicular Inputs. J. Neurosci. 32, 17620–17631 (2012).

63. Eichenbaum, H. & Lipton, P. A. Towards a functional organization of the medial temporal lobe memory system: Role of the parahippocampal and medial entorhinal cortical areas. Hippocampus 18, 1314–1324 (2008).

64. Loring, D. W. et al. The Rey Auditory Verbal Learning Test: Cross-validation of Mayo Normative Studies (MNS) demographically corrected norms with confidence interval estimates. J. Int. Neuropsychol. Soc. 29, 397–405 (2023).

65. Tulving E. (1972). Episodic and Semantic Memory. In E. Tulving, & W. Donaldson (Eds.), Organization of Memory (pp. 381–403). Cambridge, MA Academic Press. - References - Scientific Research Publishing. https://www.scirp.org/reference/referencespapers?referenceid=2919588.

66. Nilsson, J. et al. Cerebrospinal fluid biomarker panel for synaptic dysfunction in a broad spectrum of neurodegenerative diseases. Brain 147, 2414–2427 (2024).

67. Carson, R. E. et al. Imaging of Synaptic Density in Neurodegenerative Disorders. J. Nucl. Med. Off. Publ. Soc. Nucl. Med. 63, 60S–67S (2022).

68. Palombo, M. et al. SANDI: A compartment-based model for non-invasive apparent soma and neurite imaging by diffusion MRI. NeuroImage 215, 116835 (2020).

69. Behrenbruch, N. et al. A physically and mentally active lifestyle relates to younger brain and cognitive age. GeroScience 48, 1853–1873 (2026).

70. Schmid, N. S., Ehrensperger, M. M., Berres, M., Beck, I. R. & Monsch, A. U. The Extension of the German CERAD Neuropsychological Assessment Battery with Tests Assessing Subcortical, Executive and Frontal Functions Improves Accuracy in Dementia Diagnosis. Dement. Geriatr. Cogn. Disord. Extra 4, 322–334 (2014).

71. Carson, N., Leach, L. & Murphy, K. J. A re-examination of Montreal Cognitive Assessment (MoCA) cutoff scores. Int. J. Geriatr. Psychiatry 33, 379–388 (2018).

72. Behrenbruch, N. et al. A physically and mentally active lifestyle relates to younger brain and cognitive age. GeroScience 48, 1853–1873 (2026).

73. Olsen, R. K. et al. Human anterolateral entorhinal cortex volumes are associated with cognitive decline in aging prior to clinical diagnosis. Neurobiol. Aging 57, 195–205 (2017).

74. SPM12 Software - Statistical Parametric Mapping. https://www.fil.ion.ucl.ac.uk/spm/software/spm12/.

75. Behzadi, Y., Restom, K., Liau, J. & Liu, T. T. A component based noise correction method (CompCor) for BOLD and perfusion based fMRI. NeuroImage 37, 90–101 (2007).

76. Kriegeskorte, N., Mur, M. & Bandettini, P. A. Representational similarity analysis - connecting the branches of systems neuroscience. Front. Syst. Neurosci. 2, (2008).

77. Ichise, M. et al. Linearized Reference Tissue Parametric Imaging Methods: Application to [11C]DASB Positron Emission Tomography Studies of the Serotonin Transporter in Human Brain. J. Cereb. Blood Flow Metab. 23, 1096–1112 (2003).

78. Desikan, R. S. et al. An automated labeling system for subdividing the human cerebral cortex on MRI scans into gyral based regions of interest. NeuroImage 31, 968–980 (2006).

79. Mormino, E. C. et al. Tau PET imaging with 18F-PI-2620 in aging and neurodegenerative diseases. Eur. J. Nucl. Med. Mol. Imaging 48, 2233–2244 (2021).

80. Glasser, M. F. & Essen, D. C. V. Mapping Human Cortical Areas In Vivo Based on Myelin Content as Revealed by T1- and T2-Weighted MRI. J. Neurosci. 31, 11597–11616 (2011).

