## Supplemental Materials for "Entorhinal grid coding as a functional link between tau accumulation and episodic memory in human aging"

**Supplementary Fig. S1 | Right entorhinal grid-cell-like activity does not predict episodic memory performance.**

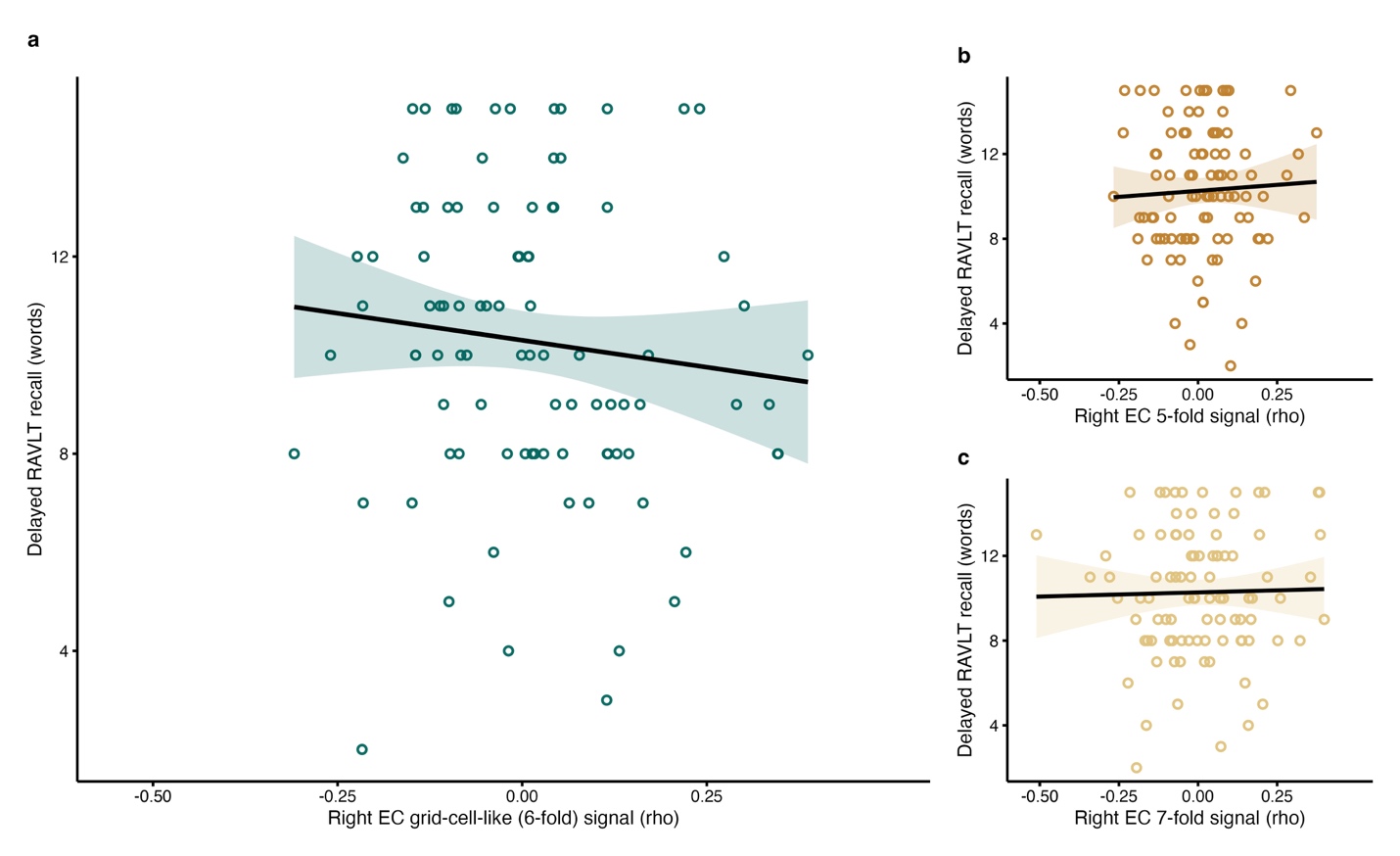

**a** Association between right entorhinal 6-fold grid-cell-like activity and delayed RAVLT recall, adjusted for age. Right entorhinal 6-fold grid-cell-like activity did not predict delayed RAVLT recall beyond age (standardized β = −0.10, 95% credible interval = −0.30 to 0.10, posterior probability of direction = 0.853; BF₁₀ = 0.43). **b,c**: Control analyses testing whether delayed RAVLT recall was associated with right entorhinal non-grid periodicities. The corresponding (**b**) 5-fold (standardized β = 0.05, 95% credible interval = −0.16 to 0.25, posterior probability of direction = 0.68) and (**c**) 7-fold (standardized β = 0.02, 95% credible interval = −0.18 to 0.22, posterior probability of direction = 0.56) control analyses are shown for comparison. Points show raw delayed-recall scores and raw grid-cell-like signal values. Regression lines show the model-estimated association adjusted for age, with age fixed at the sample mean.

**Supplementary Fig. S2 | Structural and microstructural measures do not explain left entorhinal grid-cell-like activity.**

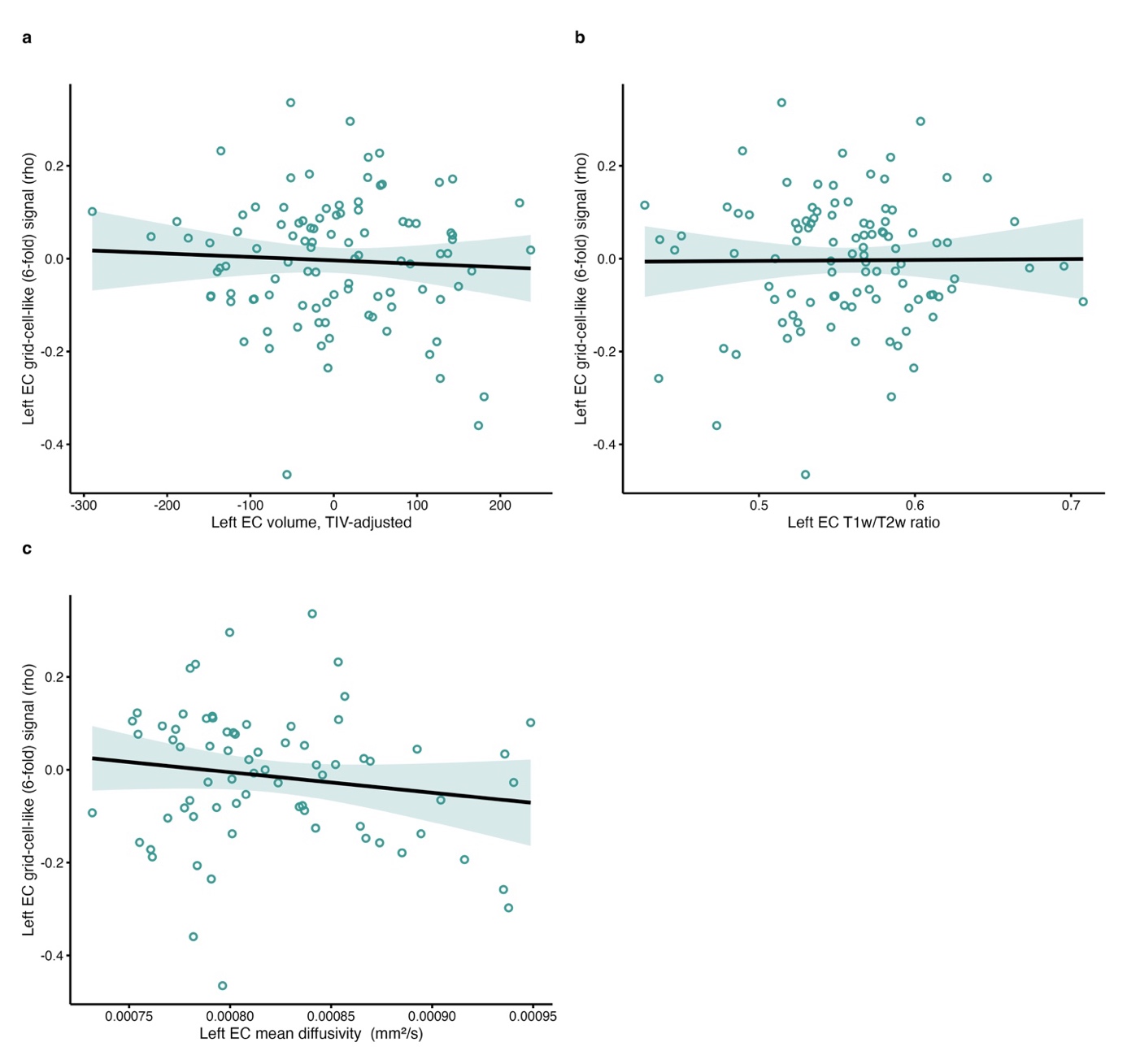

**a–c** Associations between left entorhinal 6-fold grid-cell-like activity and left entorhinal structural or microstructural measures, adjusted for age. Left entorhinal 6-fold grid-cell-like activity was not associated with TIV-adjusted left entorhinal volume (**a**; standardized β = −0.05, 95% credible interval = −0.26 to 0.15, posterior probability of direction = 0.692; BF₁₀ = 0.34), left entorhinal T1w/T2w ratio (**b**; standardized β = 0.01, 95% credible interval = −0.21 to 0.22, posterior probability of direction = 0.525; BF₁₀ = 0.31), or left entorhinal mean diffusivity (**c**; standardized β = −0.15, 95% credible interval = −0.39 to 0.09, posterior probability of direction = 0.899; BF₁₀ = 0.48). Together, these analyses suggest that individual differences in left entorhinal grid-cell-like activity were not explained by macroscopic structural variation or microstructural tissue changes in the same region. Points show individual participants. Regression lines show the model-estimated association adjusted for age, with 95% confidence intervals. *BF₁₀ values compare models including each structural or microstructural measure plus age against age-only models.*

**Supplementary Fig. S3 | Left entorhinal tau burden and temporal stability of neural representations.**

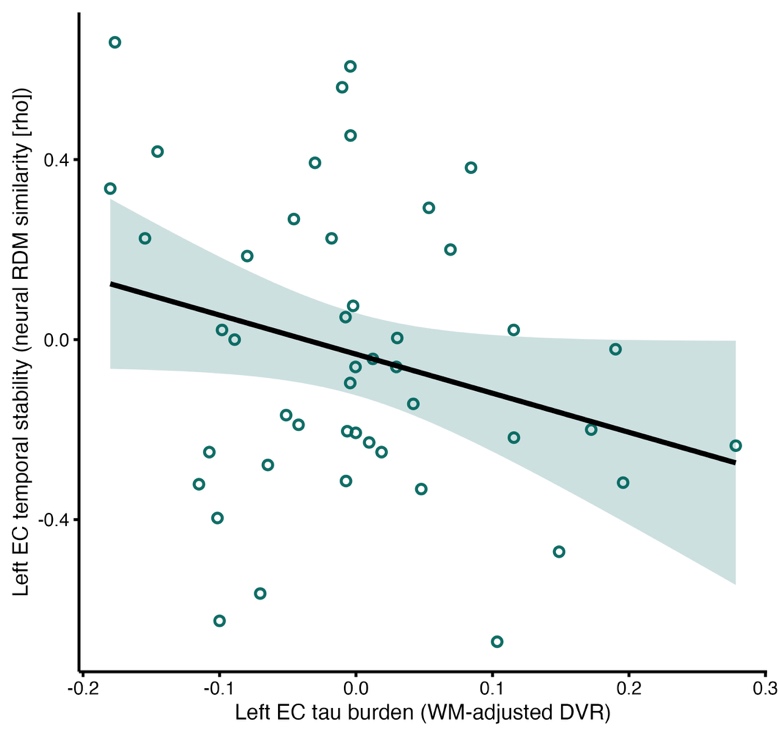

Association between left entorhinal cortex (EC) tau burden and temporal stability of left EC neural representations, adjusted for age. Higher left EC tau burden showed a negative association with temporal stability of left EC activity patterns, indexed by representational dissimilarity matrix similarity, although this effect was not reliable because the 95% credible interval overlapped zero (standardized β =-0.26 95% CrI [−0.55, 0.02], pd = 0.97). Points show individual participants. Regression lines show the model-estimated association adjusted for age with 95% confidence interval. Left EC tau burden is residualised for local white matter binding.

**Supplementary Table 1 | Sample characteristics**

| **Characteristic** | **fMRI** | **fMRI+VLMT** | **fMRI + DTI** | **fMRI + PET** |
| --- | --- | --- | --- | --- |
| Participants, n | 100 | 93 | 72 | 46 |
| Female, n (%) | 50 (50.0%) | 45 (48.4%) | 35 (48.6%) | 18 (39.1%) |
| Age, years | 76.34 (7.67) | 76.29 (7.74) | 75.56 (7.61) | 76.28 (7.67) |
| APOE ε4 carrier, n (%) | 14 (19.7%) | 13 (18.6%) | 13 (20.3%) | 6 (14.3%) |
| Aβ42/40 ratio | 0.087 (0.012) | 0.087 (0.011) | 0.087 (0.013) | 0.088 (0.011) |
| MMSE score | 28.83 (1.05) | 28.82 (1.05) | 28.90 (1.08) | 28.59 (1.07) |
| CERAD-Plus global z-score | 0.43 (0.48) | 0.44 (0.47) | 0.50 (0.44) | 0.45 (0.42) |
| VLMT delayed recall (n words) | — | 10.28 (3.02) | 10.46 (2.98) | 10.22 (3.28) |
| MTL tau burden | — | — | — | 0.01 (0.036) |
| Left entorhinal tau burden | — | — | — | -0.01 (0.101) |

Values are shown for the full fMRI sample and for the modality-specific subsamples used in the main analyses. The fMRI sample was used for analyses of entorhinal volume and T1w/T2w-derived intracortical tissue contrast. The fMRI + VLMT subsample was used to test the association between entorhinal grid-cell-like signal and episodic memory performance. The fMRI + DTI subsample was used for analyses relating grid-cell-like signal to diffusion-derived mean diffusivity. The fMRI + PET subsample was used for analyses relating tau PET burden to grid-cell-like signal and for the mediation analysis linking medial temporal tau burden, left entorhinal grid-cell-like signal, and delayed word-list recall. Values are presented as mean (SD) unless otherwise indicated. APOE ε4 carrier status is shown as n (%). Aβ42/40 ratio refers to plasma Aβ42/40. MTL and left entorhinal tau burden are shown as white-matter-adjusted DVR residuals. Dashes indicate measures not available or not applicable for that subsample.

**Supplementary Table 2 | Control bayesian mediation analysis**

| **Analysis** | **Effect** | **Path** | **Standardized beta (95% CrI)** |
| --- | --- | --- | --- |
| **Sensitivity analysis**: unadjusted MTL DVR | a-path | MTL tau → left EC grid-cell-like signal | −0.30 [−0.59, −0.01] |
| **Sensitivity analysis**: unadjusted MTL DVR | b-path | left EC grid-cell-like signal → delayed word-list recall | 0.30 [0.03, 0.56] |
| **Sensitivity analysis**: unadjusted MTL DVR | Direct effect | MTL tau → delayed word-list recall | −0.25 [−0.51, 0.02] |
| **Sensitivity analysis**: unadjusted MTL DVR | Indirect effect | MTL tau → left EC grid-cell-like signal → delayed word-list recall | −0.09 [−0.24, 0.005] |
| **EC mediation**: WM-adjusted DVR | a-path | left EC tau → left EC grid-cell-like signal | −0.46 [−0.73, −0.19] |
| **EC mediation**: WM-adjusted DVR | b-path | left EC grid-cell-like signal → delayed word-list recall | 0.28 [−0.01, 0.57] |
| **EC mediation**: WM-adjusted DVR | Direct effect | left EC tau → delayed word-list recall | −0.19 [−0.48, 0.10] |
| **EC mediation**: WM-adjusted DVR | Indirect effect | left EC tau → left EC grid-cell-like signal → delayed word-list recall | −0.13 [−0.31, 0.003] |

Note. All effects are standardized Bayesian estimates from the mediation model. Values in brackets indicate 95% credible intervals (CrI); MTL = medial temporal lobe; EC = entorhinal cortex.
